# CTP synthase forms cytoophidia in archaea

**DOI:** 10.1101/830091

**Authors:** Shuang Zhou, Hua Xiang, Ji-Long Liu

## Abstract

CTP synthase (CTPS) is an important metabolic enzyme that catalyzes the rate-limiting reaction of de novo synthesis of the nucleotide CTP. Since 2010, a series of studies have demonstrated that CTPS can form filamentous structures termed cytoophidia in bacteria and eukaryotes. However, it remains unknown whether cytoophidia exist in archaea, the third domain of life. Using *Haloarcula hispanica* as a model system, here we demonstrate that CTPS forms distinct intracellular compartments in archaeal cells. Under stimulated emission depletion (STED) microscopy, we find that some HhCTPS compartments have elongated filamentous structures, resembling cytoophidia in bacteria and eukaryotes. When *Haloarcula* cells are cultured in low-salt medium, the occurrence of cytoophidia increases dramatically. Moreover, overexpression of CTPS or glutamine analog treatment promotes cytoophidium assembly in *H. hispanica.* Our study reveals that CTPS forms cytoophidia in all three domains of life, suggesting that this is an ancient property of CTPS.

## Introduction

CTP (cytidine triphosphate) is the basic building block of RNA and DNA and the key precursor in the biosynthesis of membrane phospholipids (Carman and Henry, 1989). The synthesis of CTP is the last committed step in pyrimidine nucleotide de novo synthesis catalyzed by CTP synthase (CTPS) (Koshland Jr and Levitzki, 1974). This catalytic reaction includes the ATP-dependent phosphorylation at the C-4 position of UTP to form intermediate 4-phosphoryl UTP, which is reacted with ammonia from glutamine hydrolysis to generate CTP (Lieberman, 1956; Long and Koshland, 1978). GTP activates glutamine hydrolysis by allosterically binding to CTPS to form a covalent glutaminyl enzyme intermediate (Levitzki and Koshland, 1972; Long and Pardee, 1967). The product CTP is a competitive inhibitor of the enzyme, which is critical to maintain the intracellular CTP level (Endrizzi et al., 2005; Ostrander et al., 1998).

Compartmentalization is the process of formation of cellular compartments which play a key role in homeostasis (Liu, 2016). Compartments are often defined as membrane-enclosed regions such as mitochondria and chloroplasts (Diekmann and Pereira-Leal, 2013), while also including protein-based membraneless organelles such as purinosomes (An et al., 2008) and Cajal bodies (Cajal, 1903). In 2010, CTPS was found to compartmentalize into filamentous structures in *Drosophila*. These structures are termed cytoophidia (cellular snakes in Greek) (Liu, 2010). Since then, independent studies have discovered that CTPS can form cytoophidia in bacteria (*E. coli* and *Caulobacter crescentus*) (Ingerson-Mahar et al., 2010), and eukaryotes such as budding yeast, fission yeast, human cells and plant cells (Carcamo et al., 2011; Chen et al., 2011; Daumann et al., 2018; Ingerson-Mahar et al., 2010; Noree et al., 2010; Zhang et al., 2014). The phenomenon that CTPS forms cytoophidia is conserved across bacteria and eukaryotes.

Life can be divided into three domains: archaea, bacteria and eukarya (Woese et al., 1990). Several studies have found evidence of CTPS forming cytoophidia in bacteria and eukaryotes, which raises a very interesting question: we would like to know if this process occurs in archaea. The reason for this interest is that many archaea survive in extreme environments in which most microorganisms are not able to survive. So we want to ask the following three questions: 1) Can CTPS form intracellular structures in archaea? 2) Under what conditions can the intracellular structure be formed? 3) What is the relationship between the formation of intracellular structures and the physiology of archaea?

To answer those questions, we chose the halophilic archaeon *Haloarcula hispanica (H. hispanica)* ATCC 33960 as a model system. *Haloarcula species* are extremophilic halophiles requiring 1.5 M to saturated salt concentration for growth (Madern et al., 2000; Thombre et al., 2016). *H. hispanica* was isolated from a solar saltern in Spain, and is a classic model in archaeal research (Juez et al., 1986). *H. hispanica* can survive well in a medium containing optimal NaCl concentration at 3.5-4.2 M. It grows slowly in the case of low salt. Other conditions are also important for *H. hispanica* growth; for example, it requires at least 0.005 M Mg^2+^ concentration. In addition, the optimum growth temperature is 35-40°C and the optimal pH is 7.0 (Garrity, 2012). We found that there were few cytoophidia (about 1%) under normal growing conditions (AS168 medium, 37 ℃). Furthermore, growth and stress regulated *Haloarcula hispanica* CTPS cytoophidia formation.

## Results

### CTPS forms cytoophidia in *H. hispanica*

CTPS is evolutionarily conserved among all three domains of life: archaea, bacteria and eukarya (**Figure 1**). The *H. hispanica* CTP synthase (HhCTPS) gene is located in Chromosome I (Liu et al., 2011b), and HhCTPS contains two conserved regions: N terminal (residues 16 to 277, ligase domain) and C terminal (residue 303 to 535, glutaminase, type 1 glutamine amidotransferase). We tried to align HhCTPS with some classical cytoophidium-forming CTPS types and well-studied crystal structures of CTPS. There is a significant similarity between CTP synthase from *H. hispanica* and proteins from *Thermus thermophiles*, humans, *Saccharolobus solfataricus*, *Drosophila*, *E. coli*, *Mycobacterium tuberculosis* and *Saccharomyces cerevisiae*, which share 46% - 53% sequence identity **(Figure S1; Table S1)**.

**Figure 1.**
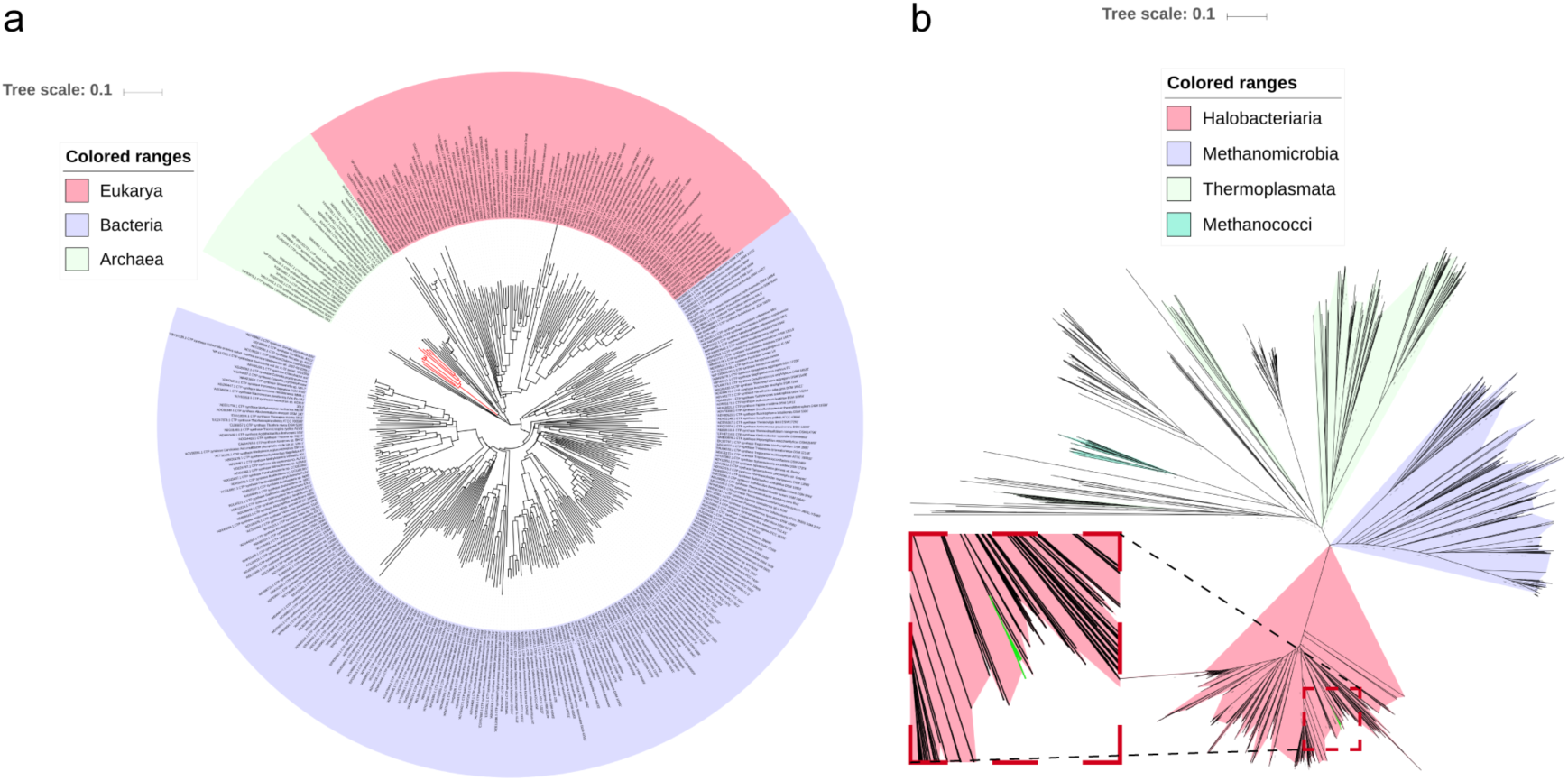
Neighbor-Joining (Saitou and Nei, 1987) phylogenetic trees of CTP synthase proteins. **a,** Representative CTPS proteins from the three life domains are selected and aligned for creating the phylogram. The CTPS proteins from eukarya, bacteria and archaea are respectively shaded in pink, lilac, pale green. Proteins from halophiles are indicated in red. **b**, Almost all CTPS proteins in Euryarchaeota were used for building the phylogenetic tree, which contains clusters from Halobacteria (pink), Methanomicrobia (lilac), Thermoplasmata (pale green), Methanococci (cyan). The *Haloarcula* proteins are indicated in green which is shown in the red dotted box.

In order to visualize the *H. hispanica* CTPS in *H. haloarcula*, we fused modified pSMRSGFP to the C-terminus of CTPS and visualized the distribution of HhCTPS-GFP (pSMRSGFP knock-in strain) under laser scanning confocal microscopy and super-resolution microscopy (**Figure 2a-c**). We found that CTPS is mostly diffusely distributed under normal conditions. Only a small number of cells can form foci (**Figure 2b**). The intracellular compartmentalized foci were relatively small and it was difficult to determine the shape of those foci under conventional confocal microscopy.

**Figure 2.**
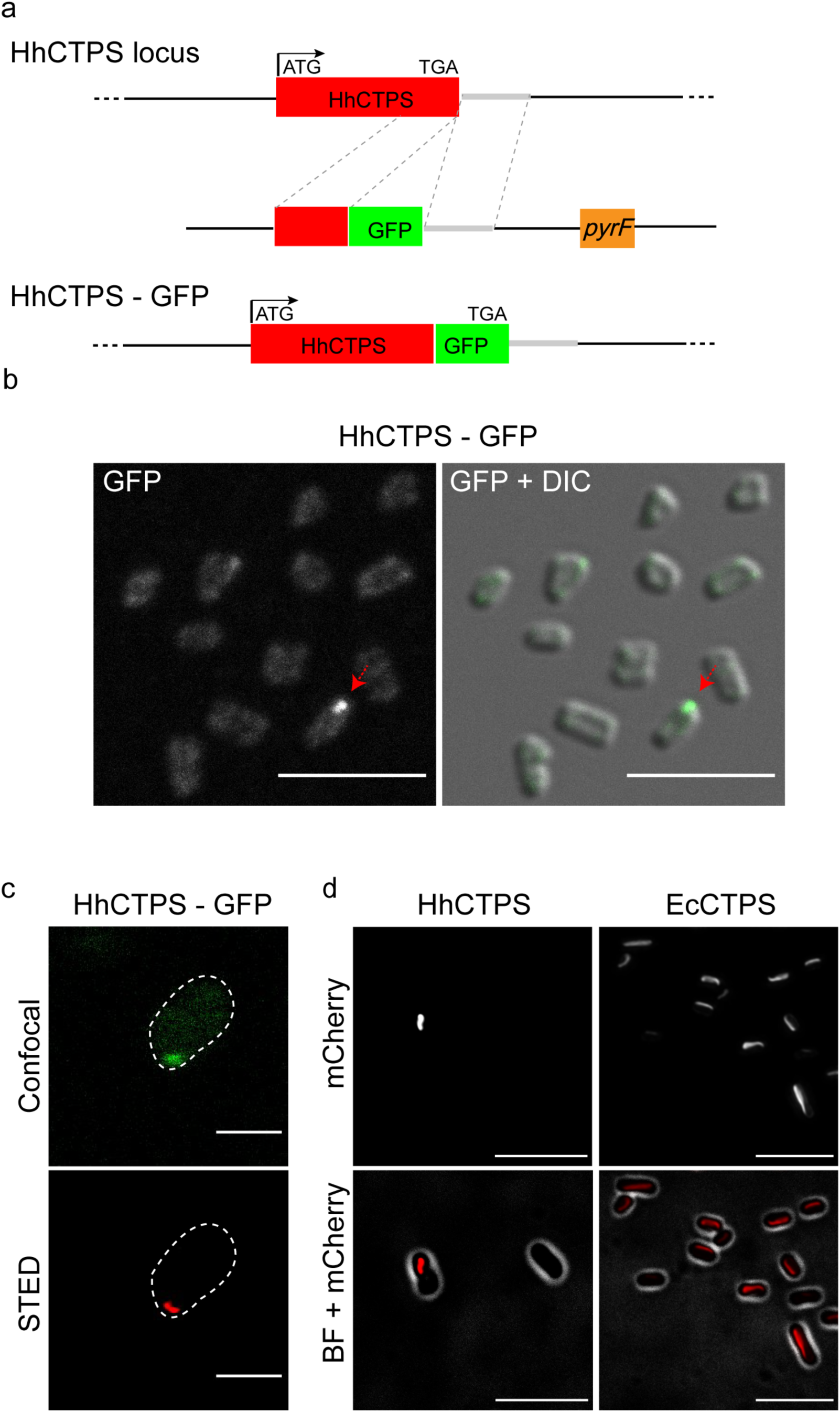
CTP synthase forms cytoophidia in *H. hispanica*. **a**, C-terminal targeting of endogenous *H. hispanica* CTPS (HhCTPS) with GFP. **b**, Confocal images of *H. hispanica* show the compartmentalized capability of HhCTPS in the late log phase. Arrows point to the punctate structure. The DIC image represents the cell contour. **c**, In *H. hispanica* cell, a compartmented CTPS structure (green, upper pane) observed under conventional confocal microscopy shows an elongated shape (red, lower panel) under stimulated emission depletion (STED) microscopy. This elongated filamentous structure in the archaeal cell resembles cytoophidia described in bacteria and eukaryotes. **d**, Heterologous expression of HhCTPS in *E. coli.* Homogenous *E. coli* CTPS (EcCTPS) forms longer cytoophidia in more *E. coli* cells than heterogeneous HhCTPS. Both HhCTPS and EcCTPS are tagged with mCherry. *H. hispanica* is collected in the late log phase, *E. coli* is collected in stationary phase. Scale bars, 5 μm in (b) and (d), 1 μm in (c).

We also wanted to see how the intracellular compartmentalized structures in *H. hispanica* differ from cytoophidia in bacteria and eukaryotes. Specifically, we wished to know whether the HhCTPS foci are completely circular or elongated. Under stimulated emission depletion (STED) microscopy, we observed that some HhCTPS-forming compartmentalized structures that look like a rod or dot under confocal are actually lengthened structures (**Figure 2c**). What’s more, we quantified the endogenous cytoophidia using STED images of *H. hispanica* and confocol images of the other species including *Homo sapiens* (Sun and Liu, 2019), *Drosophila melanogaster, Schizosaccharomyces pombe*, *Saccharomyces cerevisiae* **(Figure S2)**. We found that the axia ratios of *H. hispanica* and *S. pombe* are smaller than for the other species, mainly concentrated around 1.5 **(Figure S2)**. Our results indicate that HhCTPS can form elongated filamentous structures in *H. hispanica*. Therefore, we concluded that cytoophidia do exist in this species.

### *H. hispanica* CTPS forms cytoophidia in *E. coli*

To determine whether HhCTPS can form cytoophidia in other species independent of its own intracellular environment, we expressed mCherry-labeled HhCTPS in *E. coli*. We found that HhCTPS can form cytoophidia in *E. coli* (**Figure 2d**), but they are somewhat different to those formed by the *E. coli* CTPS (EcCTPS). In *E. coli* cells, homogenous EcCTPS forms longer cytoophidia in more cells than heterogenous HhCTPS. The results demonstrate that Haloarcula CTPS has the ability to form cytoophidia in *E. coli*.

### A glutamine analog promotes cytoophidium assembly in *H. hispanica*

Eukaryotic CTPS uses glutamine as a substrate. It was found that 6-diazo-5-oxo-l-norleucine (DON), an analog of glutamine, inhibits CTPS activity by irreversibly binding to its glutamine amidotransferase domain. Meanwhile, DON promotes cytoophidium assembly in *Drosophila* and human cells (Boeke et al., 1984; Levitzki and Koshland Jr, 1971), but inhibits the formation of cytoophidia in bacteria (Ingerson-Mahar et al., 2010). To understand how DON affects HhCTPS in *H. hispanica*, cells were cultured in medium supplemented with DON. We observed that DON treatment induces cytoophidium assembly in *H. hispanica* cells **(Figure S3)**. Meanwhile, DON treatment inhibits cell growth in a concentration-dependent manner **(Figure S3b)**. Our data indicate that the effect of DON on HhCTPS-forming cytoophidia is similar to that on *Drosophila* and human CTPS, but different to that on bacterial CTPS.

### Low salinity induces cytoophidium formation in *H. hispanica*

Next, we wanted to know how formation of cytoophidia is related to the physiological characteristics of *H. hispanica*. We cultured *H. hispanica* under different conditions. We found that the *H. hispanica* cells cultured in low-salt medium (1.5 M NaCl AS168) become round (like a balloon) from the original polymorphism (rod, circle, square, triangle, irregular shapes) due to osmotic pressure (**Figure 3a-d**). In comparison, the cell shape became bigger and longer under DON treatment (**Figure 3**). Quantification analysis of cytoophidium locations in DON-treated cells showed that more than half of the dividing cells contain one or two cytoophidia **(Figure S4)**. It seemed that cell division was suppressed because of the inhibition of HhCTPS by DON. Then we tried to culture HhCTPS-GFP in nutrient-limited minimal medium (MG medium) (Han et al., 2009) or resuscitation medium to arrest or promote cell growth. We found that cells were also deformed into a bigger or longer shape, but the cytoophidia had no effects **(Figure S5)**.

**Figure 3.**
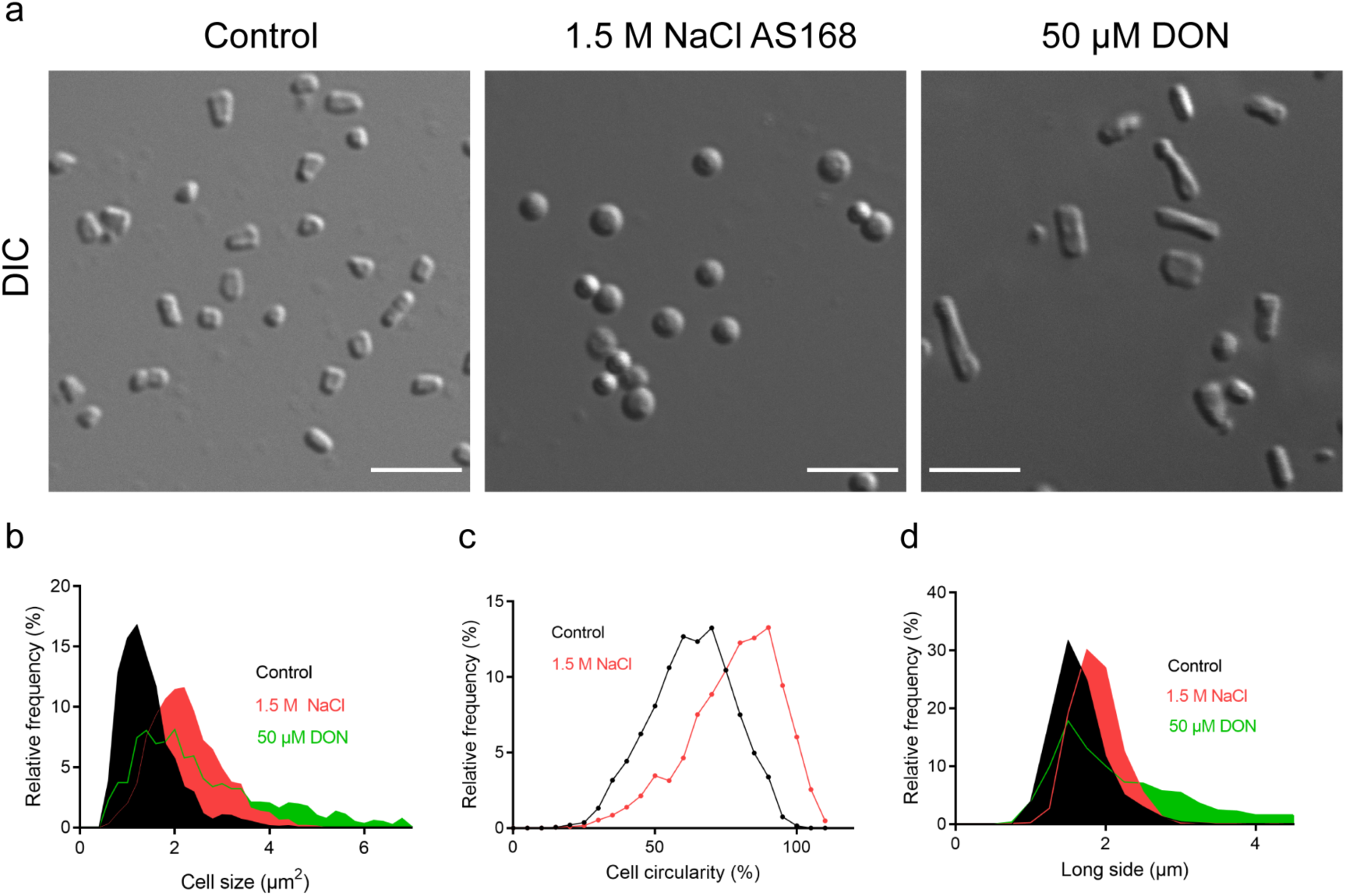
Low salinity and DON treatment change cell shapes in *H. hispanica*. **a**, DIC images of HhCTPS-GFP in normal conditions, low salinity medium, DON-treated medium, on the third day of cultivation. **b**, **c**, **d**, Images were analyzed for cell size (**b**), circularity (**c**), and length (the longer side of minimum bounding rectangle) (**d**). n = 1897 cells in normal conditions, 1181 cells in low salinity medium, and 1874 cells in DON-treated medium. Mean ± SD. Scale bars, 5 μm.

In addition, to our surprise, HhCTPS can form a large number of intracellular structures under low-salt conditions. In 1.5M NaCl AS168 medium (**Figure 4a, b**), the *H. hispanica* cells grew very slowly, with growth almost stopping (**Figure 4c, d**). After cultivation for about 3 days in this medium, HhCTPS-GFP cells were transferred to normal AS168 medium (**Figure 4e**). Cell growth continued but the abundance of HhCTPS compartmentalized structures decreased. If HhCTPS-GFP cells were maintained in the low salinity medium, there was still growth cessation with higher abundance of compartmentalized structures (**Figure 4e-h**). These results show that low salinity cannot support the normal cell growth of halophiles. Salt shortage triggers HhCTPS compartmentalization, which might be to provide a repository for enzymes to keep cells in a static state.

**Figure 4.**
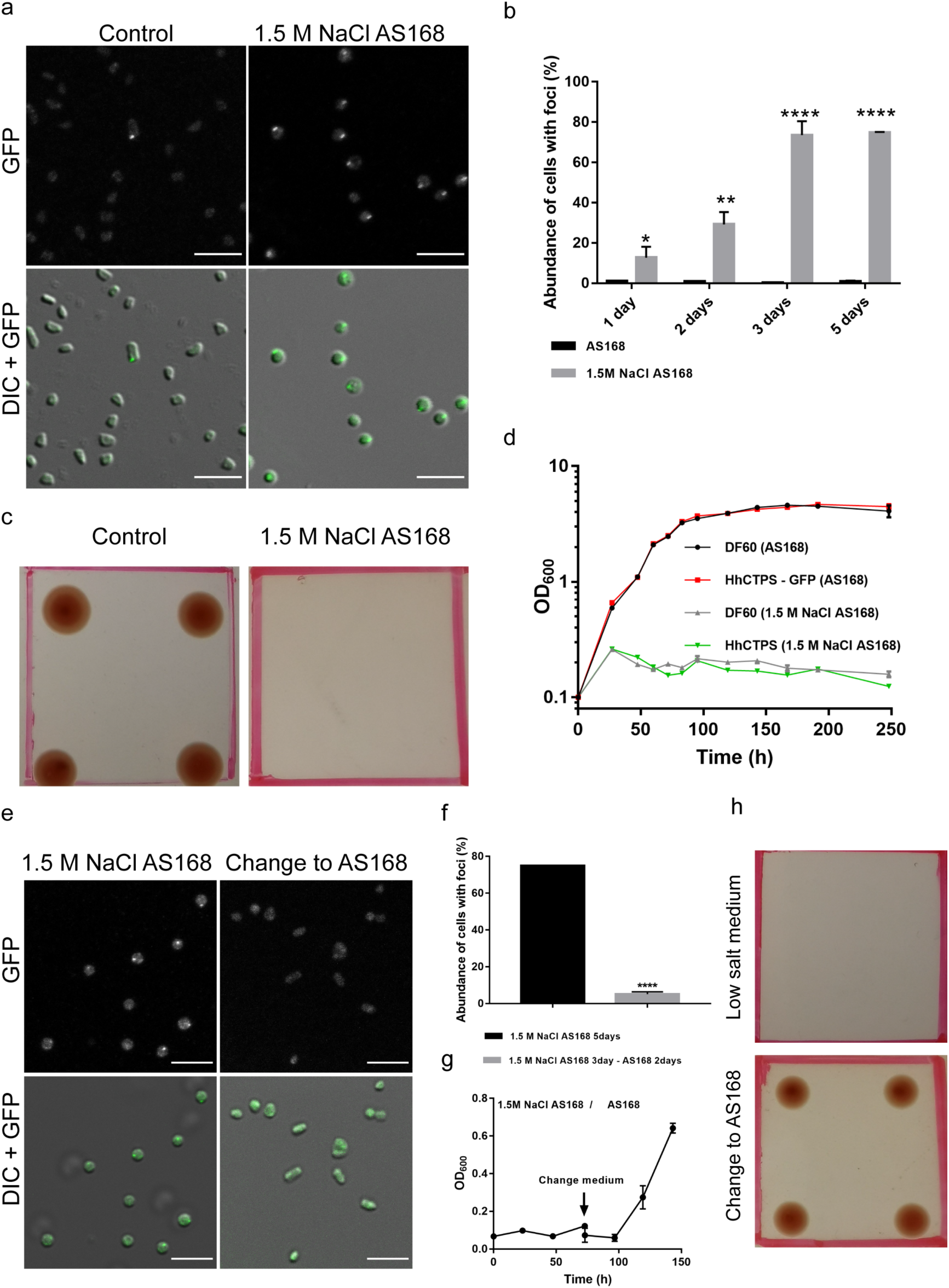
Low salinity promotes the compartmentation of CTP synthase *H. hispanica*. **a**, Many compartmentalized structures appear in the low salinity medium. **b**, Compartmentation ratio increases in the low salt medium. Cells were collected at the first day (n = 1117 cells in AS168, 1553 cells in 1.5 M NaCl AS168), the second day (n = 2538 cells in AS168, 2180 cells in 1.5 M NaCl AS168), the third day (n = 3069 cells in AS168, 2118 cells in 1.5 M NaCl AS168) and the fifth day (n = 2639 cells in AS168, 1001 cells in 1.5 M NaCl AS168) of cultivation. Mean ± SD, *p <0.05, **p < 0.01, ****p < 0.0001. **c**, **d**, *H. hispanica* cannot grow normally in 1.5 M NaCl AS168 medium. There are red colonies in the AS168 plate (left pane), but no colonies in 1.5 M NaCl AS168 after 10 days incubation (right pane) (**c**). The growth curve of *H. hispanica* in the two media with a log scale for y axis (**d**). **e**, **f**, **g**, **h**, Salt supplementation recovers cell growth, but decreases the abundance of punctate structures. Confocal images (**e**), ratio of compartmentalized structures (**f**) (n = 1001 cells for low salt culture, 1072 cells for normal AS168 culture), growth curve (**g**) and plates (**h**) show that salinity recovery diminishes compartmentation of HhCTPS-GFP. Positions of four repeat inoculations per plate are at the four corners of the red squares. Mean ± SD, ****p < 0.0001. Scale bars, 5μm.

STED images show elongated structure of HhCTPS cytoophidia. Using STED, we observed that the compartmentalized structures in cells cultured in low-salt medium exhibited an elongated shape **(Figure S6)**. The lengths of cytoophidia were significantly increased in DON-treated cells **(Figure S6b, c)**.

### Overexpressing *Haloarcula* CTPS promotes cytoophidium formation

We considered whether the level of CTPS has an effect on cytoophidium formation. Cytoophidia were induced in many *H. hispanica* cells when CTPS was overexpressed (**Figure 5a**). STED images showed that cytoophidia induced by HhCTPS overexpression were indeed elongated structures (**Figure 5b**). Moreover, the formation of cytoophidia changes with the growth phases of archaea (**Figure 5c,d**). There was a dramatic decrease of abundance of foci on the first day. Narrowing the time range to 0-60h then revealed an obvious drop followed by a rise in abundance **(Figure S7)**. The number of cytoophidia in the early stage of growth was reduced, but the GFP signal became stronger. The abundance of cytoophidia in the logarithmic phase and subsequent growth phases was almost unchanged, but the size of these structures decreased initially and then increased (**Figure 5e,f**).

**Figure 5.**
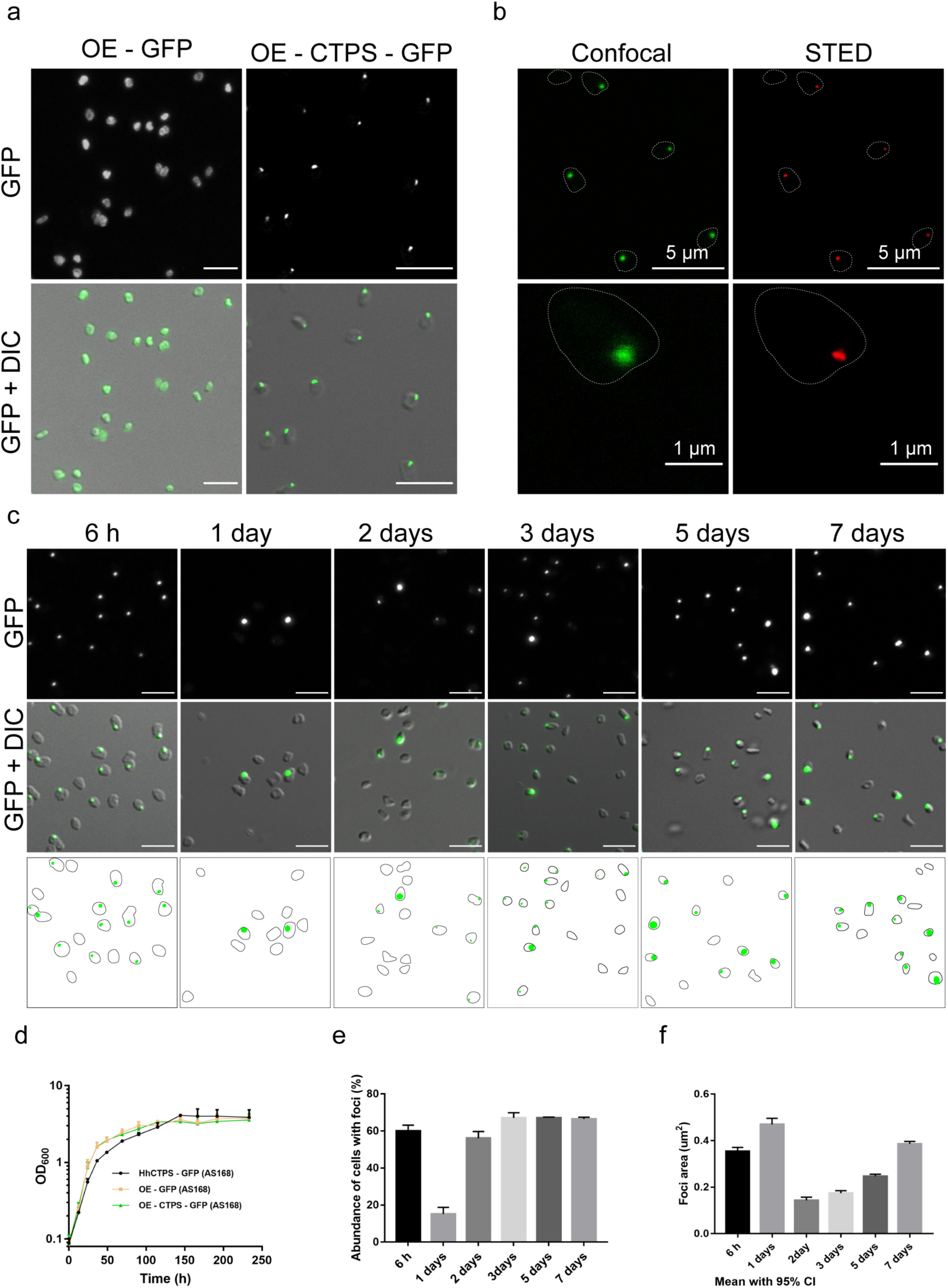
Overexpressing CTPS promotes cytoophidium assembly in *H. hispanica*. **a**, Confocal images of OE-GFP and OE-HhCTPS-GFP in late log phase in AS168 medium. **b**, STED images of OE-HhCTPS-GFP. **c**, Images of OE-HhCTPS-GFP sampled at different time points. **d**, Growth curve of DF60, OE-GFP and OE-HhCTPS-GFP with a log scale for y axis**. e**, **f**, Quantification analysis of images of OE-HhCTPS-GFP for the abundance of compartmentalized structures and foci size. Mean ± SD in (**d**), mean with 95% CI in (**e, f**). n=1210 cells for 6 h, 1247 cell for 1 day, 1378 cells for 2 days, 1540 cells for 3 days, 1961 cells for 5 days, and 1938 cells for 7 days in (**e**). n= 814 cells for 6 h, 478 for 1 day, 619 cells for 2 days, 740 cells for 3 days, 1230 cells for 5 days, and 1330 cells for 7 days in (**f**). Scale bars, 5 μm in (**a, c**).

### Yeast extract removal affects cytoophidium formation in *H. hispanica*

Low salinity (1.5 M NaCl AS168 medium) induced cytoophidium formation. We next investigated whether altering other components of the AS168 medium (per liter, 200 g NaCl, 20 g MgSO_4_·7H_2_O, 2 g KCl, 3 g trisodium citrate, 1 g sodium glutamate, 50 mg FeSO_4_·7H_2_O, 0.36 mg MnCl_2_·4H_2_O, 5 g Bacto Casamino acids, 5 g yeast extract, pH 7.2) could affect cytoophidium formation. To do this, we cultured HhCTPS-GFP cells in AS168 media, each with a different component subtracted or reduced. In addition to sodium chloride that can affect cytoophidia formation, only Mg^2+^ and yeast extract have influence on either cell shape or cytoophidia formation among these components. We found that Mg^2+^ deprivation made the cells rounder, similarly to low salt influence, but did not affect HhCTPS cytoophidium formation **(Figure S8)**. Interestingly, we found that HhCTPS also formed a number of cytoophidia (**Figure 6a,b**), noted as elongated structures under STED microscopy (**Figure 6c**), in the cells cultured in yeast extract-subtracted AS168 medium. Moreover, the growth curve of HhCTPS-GFP indicated that the growth of *H. hispanica* is inhibited in yeast extract-subtracted AS168 medium (**Figure 6d**).

**Figure 6.**
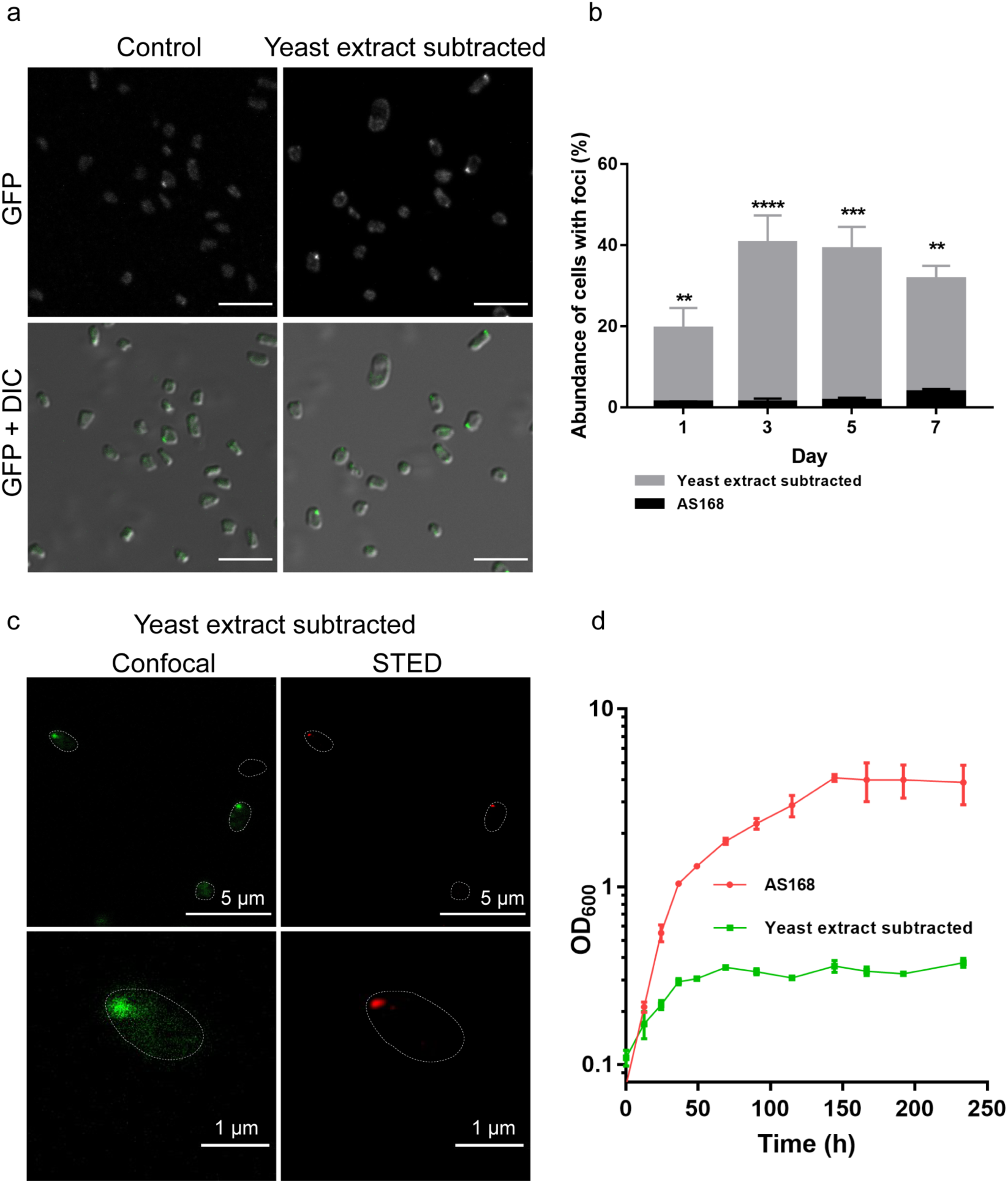
Yeast extract deprivation promotes cytoophidium formation in *H. hispanica.* **a**, Confocal images of HhCTPS-GFP in normal and yeast extract-subtracted AS168 on the third day of cultivation. **b**, Quantification of the cells containing compartmentalized structures. n = 985 cells for 1 day, 1322 cell for 3 days, 1445 cells for 5 days, and 1259 cells for 7 days in AS168. n = 1316 cells for 1 day, 1436 cells for 3 days, 1042 cells for 5 days, and 1172 cells for 7 days in yeast extract subtracted AS168. Mean ± SD. **c**, Comparison of confocal and STED images of HhCTPS-GFP in yeast extract-subtracted AS168 on the third day of cultivation. **d**, Growth curve of HhCTPS with a log scale for y axis. Scale bars, 5 μm in (**a**) and the upper panes of (**c**). 1 μm in the lower panes in (**c**).

## Discussion

In summary, we found that CTPS can form cytoophidia in *H. hispanica*. Our study demonstrates that CTPS forms cytoophidia not only in bacteria and eukaryotes but also in archaea. Cytoophidia exist in all three domains of life, suggesting that this is an ancient and fundamental property of CTPS. Furthermore, we found that environmental stress, such as low salinity or yeast extract deprivation, can promote cytoophidium assembly in *H. hispanica*.

CTPS may be mostly diffused under normal circumstances. However, in the case of low salt, CTPS can aggregate to form cytoophidia. Meanwhile, the cell shape becomes rounder, which is similar to Mg^2+^ deprivation *in H. hispanica*. Research has shown that the Na^+^/Mg^2+^ ratio has an influence on amylase activity (ENACHE et al., 2009). When we decrease the concentration of Na+ or Mg2+, is the HhCTPS activity affected and are cytoophidia formed in response? This deserves further study. Some previous *in-vitro* studies indicated that the product nucleotide-bound filament is an inactive form of CTPS in prokaryotes, but an active form of CTPS in eukaryotes. (Barry et al., 2014; Lynch et al., 2017). However, two recent studies showed that *Drosophila* CTPS and human CTPS2 can form two kinds of filament: product-induced and substrate-induced (Lynch and Kollman, 2019a, b; Zhou et al., 2019). It is therefore necessary to further define the CTPS activity state with regard to the different types of filament. We worked to purify the HhCTPS expressed by E. coli, and found that it has no enzyme activity. However, the HhCTPS protein purified from halophiles has obvious enzyme activity, which indicates that halophilic proteins need the salt environment to support normal translation or protein folding (unpublished data). We wonder which kind of filament HhCTPS can form *in vitro*: product-bound, substrate-bound, or both? In addition, do different Na^+^/Mg^2+^ ratios, or Na^+^ or K^+^ concentrations affect filament formation or enzyme activity?

Low salt induced cell shape change might be related to osmotic pressure. The amino acid composition of halophilic archaea proteins is enriched in acidic residues on the surface of the protein compared with those from non-halophilic archaea. An increase in the number of acidic residues is thought to promote the correct folding of proteins under high salt conditions (Fukuchi et al., 2003). Meanwhile, the walls are assembled by surface layer proteins that form the S layer. In most archaea (Kandler and König, 1998). The S-layer glycoproteins are also enriched in acidic residues (Eichler, 2003). Under low salt conditions, S-layer glycoproteins might lose function.

It is hard to maintain normal cell shape. Halophilic archaea can grow in environments where water activity (a_w_) is close to 0.75, but a_w_ below 0.90 has an inhibitory effect on most microorganisms (Stevenson et al., 2015). The amount of solute is related to the osmotic pressure of the microorganisms. When there is a significant imbalance in the osmotic pressure of the cells, it may lead to cell lysis, protein unfolding, and enzyme inactivation (Cheftel, 1995). Halophilic archaea overcome this problem by keeping compatible solutes such as potassium chloride (KCl) in the cell to balance osmotic pressure (Antranikian, 1998). The way halophilic archaea retains these salts is called “salt-in”. The largest component in the AS168 medium is NaCl. When we lowered the NaCl concentration in AS168, HhCTPS tended to compartmentalize. We deduced that there is no need for more acidic residues to resist extracellular salt during low salt conditions. As a result, these proteins tend to cluster together in order to hide the acidic residues.

DON treatment inhibited cell growth but promoted HhCTPS assembly. What’s more, more than half of the cytoophidia were located in the middle of the *H. hispanica* cell. It is known that the glutamine analog DON promotes CTPS cytoophidium formation in the human cell and Drosophila cell, but diminishes CTPS compartmentalization in bacteria (Chen et al., 2011; Ingerson-Mahar et al., 2010). CTPS utilizes either L-glutamine or ammonia as the source of nitrogen (Endrizzi et al., 2004; Willemoes, 2004). L-glutamine is the primary amino donor in *E. coli* as in animals, but the utilization of ammonia was stimulated by DON (Chakraborty and Hurlbert, 1961). This difference might be due to the bias of nitrogen donors while treating with DON. A study in *S. pombe* showed the asymmetric inheritance of both cytoplasmic and nuclear cytoophidia during cell division (Zhang et al., 2014). The cell growth inhibition and cell enlargement induced by DON treatment might be due to cell cycle arrest in *H. hispanica*. The *H. hispanica* cells can’t divide normally into daughter cells, which leads to larger or longer cells. Also of interest is that the CetZ proteins related to tubulin and FtsZ play a key role in controlling the archaeal cell shape. And some CetZ proteins are located at the middle of the cell during the log phase growth stage while being localized at one or both poles in the motile halo cells (Duggin et al., 2015). Many HhCTPS cytoophidia are located near the cell envelope during culture in the normal AS168 medium. DON treatment facilitates the cytoophida to gather at the middle of the cells, which indicates that cell cycle arrest stops cytoophidia from being inherited by daughter cells.

*H. hispanica* is one example of archaea. Our work shows that CTPS can form cytoophidia not only in eukaryotes and bacteria, but also in archaea. This work and studies in several laboratories (including ours) in the first decade of the cytoophidium field indicate that filament forming is an ancient and evolutionarily conservative phenomenon. In order to understand more about the role of the cytoophidia formed by CTPS in archaea, further exploration is necessary for other archaea species, such as thermophiles and methanogens.

## Material and Methods

### Strains and growth conditions

The strains used in this study are listed in **Supplementary Table S2**. *Escherichia coli* was grown in Luria-Bertani medium at 37°C (CHONG, 2001), with ampicillin at a final concentration of 100 μg/ml. The uracil auxotrophic (pyrF deleted) strain DF60 (Liu et al., 2011a) and its derivatives were cultured at 37°C in AS168 medium (per liter, 200 g NaCl, 20 g MgSO_4_·7H_2_O, 2 g KCl, 3 g trisodium citrate, 1 g sodium glutamate, 50 mg FeSO_4_·7H_2_O, 0.36 mg MnCl_2_·4H_2_O, 5 g Bacto Casamino acids, 5 g yeast extract, pH 7.2). For individual experiments, uracil was added at a final concentration of 50 mg/l (Cai et al., 2012). In some experiments, 1.5M NaCl AS168 medium (per liter, 88g NaCl, other components the same) was used. For DON treatments, 1 – 146 μM DON was added in the AS168 medium. For solid medium, 1.5% agar was added. Strains transformed with a suicide plasmid were cultured in yeast extract-subtracted AS168 (Dyall-Smith, 2008).

### Plasmid construction, transformation, and gene mutants

The plasmids used in this study are listed in **Supplementary Table S2** and primers are listed in **Supplementary Table S3**. For knock-in of pSMRSGFP in *H. hispanica* ATCC 33960, a 729-bp DNA fragment of pSMRSGFP gene with linker (AGSAA) (Schmidt and Pfeifer, 2013), a 663-bp DNA fragment at the 3′ region of the HhCTPS gene containing a BamH I site at the 5′ end, and a 673-bp DNA fragment located downstream of the HhCTPS operon containing a Kpn I site at the 3′ end were amplified by primer pairs HhCTPS-C-GFP-F / HhCTPS-C-GFP-R, HhCTPS-C-UF/ HhCTPS-C-UR and HhCTPS-C-DF/ HhCTPS-C-DR (**Supplementary Table S3**) from plasmid pWL502-GFP and genome. These three DNA fragments were linked and inserted into the suicide plasmid pHAR (predigested with BamHI and KpnI), and transferred into DF60 cells to knock in PSMRSGFP by the pop-in/pop-out method described previously (Liu et al., 2011a). The expression plasmid pWL502 (Cai et al., 2012) was used to overexpress HhCTPS-GFP. A DNA fragment of HhCTPS with PSMRSGFP was amplified from HhCTPS-GFP knock-in strain containing a BamH I site at 5′ end and a Kpn I site at 3′ end. This DNA fragment was inserted into plasmid pWL502 (predigested with BamHI and KpnI) and transferred into DF60 cells. pWL502-GFP (pWL502-pSMRSGFP) was used as control. All transformants were screened by PCR verification and confirmed by sequence analysis.

### Heterogeneous expression

For heterogeneous expression of HhCTPS in *E. coli*, DNA fragments of HhCTPS or EcCTPS operon with linker (GGGS) and mCherry gene were amplified by primer pairs ECCTPS-F / ECCTPS-R, mCherry-F / mCherry-R. Two DNA fragments were linked and inserted into plasmid pET28a, and transferred into Transetta (TRANSGEN BIOTECH). IPTG with a final concentration of 1 μM was added when needed.

### Preparation of fixed *H. hispanica*

Paraformaldehyde containing 20% NaCl was added to *H. hispanica* cell culture to give a final concentration of PFA of 4%. After fixing for 10 min at room temperature, cells were spun down for 3 min at 6000 rpm/min, then washed once in basal salt solution (BSS; per liter, 200 g NaCl, 20 g MgSO_4_ · 7H_2_O, 2 g KCl, and 3 g trisodium citrate). Samples were stored in BSS at 4°C.

### Light microscopy

Cell suspension (2 μl) was mixed with 2 μl 1% low melting temperature agarose containing 18% Salt Water to avoid movement of the cells during imaging (Duggin et al., 2015; Dyall-Smith, 2008). Images were acquired under 63x objectives with differential interference contrast (DIC) on a laser scanning confocal microscope (Leica SP8, Leica Microsystems CMS GmbH, Mannheim, Germany). For super-resolution imaging, the stimulated emission depletion (STED) module (Leica TCS SP8 STED 3X) was used. STED with GFP was performed with the 592 nm continuous wave (CW). Images were captured using a 100x 1.4 NA oil objective. Instrument aberration and blurring were corrected with post-acquisition deconvolution using Huygens Professional software. DIC and GFP epifluorescence images were analyzed using Fiji with the ImageJ Shape Filter Plugin. Area (size): the area enclosed by the outer contour of an object. Long side: the longer side of the minimum bounding rectangle. Circularity = *4π × [Area] / [Perimeter]* with a value of 1.0 indicating a perfect circle. As the value approaches 0.0, it indicates an increasingly elongated shape. Axial ratio = Major axis / Minor axis.

### Growth analysis

*H. hispanica* cells were diluted in growth medium to a start optical density of approximately 0.1 at 600 nm (OD_600nm_) (with growth medium used for blank). 5 ml cell cultures were incubated in 12 ml flasks or 20 ml cell cultures were incubated in 100 ml flasks at 37°C with shaking (220 rpm). The OD_600nm_ was measured for three to ten days in each flask. The growth curves represent the average of three independent samples. The cells grown in 1.5 M NaCl AS168 were centrifuged for 3 min at 6000 rpm/min and the supernatant was removed. Then the cells were washed once with AS168 or 1.5 M NaCl AS168, and finally transferred to AS168 or 1.5 M NaCl AS168 for continuous cultivation. The OD_600nm_ was measured every day in each flask. For cell growth on the agar plate, 2 μl of culture in the growth medium was inoculated onto the subsurface of another agar plate, with up to four inoculations per plate from the same liquid culture. Plates were incubated in a closed plastic box at 42°C for 10 days, and then images of the plates were acquired using a camera.

## Acknowledgments

We thank Junyu Chen, Luyao Gong and Xiaoming Li for technical assistance and the Molecular Imaging Core Facility at School of Life Science and Technology, ShanghaiTech University for providing technical support. Meanwhile, we thank Shanshan Zhang, Yuanbing Zhang for confocol images of cytoophidia from other species.

## References

An, S., Kumar, R., Sheets, E.D., Benkovic, S.J., 2008. Reversible compartmentalization of de novo purine biosynthetic complexes in living cells. Science 320, 103–106.

Antranikian, G., 1998. Biotechnology of extremophiles. Springer.

Barry, R.M., Bitbol, A.F., Lorestani, A., Charles, E.J., Habrian, C.H., Hansen, J.M., Li, H.J., Baldwin, E.P., Wingreen, N.S., Kollman, J.M., Gitai, Z., 2014. Large-scale filament formation inhibits the activity of CTP synthetase. Elife 3, e03638.

Boeke, J.D., La Croute, F., Fink, G.R., 1984. A positive selection for mutants lacking orotidine-5′-phosphate decarboxylase activity in yeast: 5-fluoro-orotic acid resistance. Molecular and General Genetics MGG 197, 345–346.

Cai, S., Cai, L., Liu, H., Liu, X., Han, J., Zhou, J., Xiang, H., 2012. Identification of the haloarchaeal phasin (PhaP) that functions in polyhydroxyalkanoate accumulation and granule formation in Haloferax mediterranei. Appl Environ Microbiol 78, 1946–1952.

Cajal, S.R., 1903. Un sencillo metodo de coloracion seletiva del reticulo protoplasmatico y sus efectos en los diversos organos nerviosos de vertebrados e invertebrados. Trab. Lab. Invest. Biol.(Madrid) 2, 129–221.

Carcamo, W.C., Satoh, M., Kasahara, H., Terada, N., Hamazaki, T., Chan, J.Y., Yao, B., Tamayo, S., Covini, G., von Muhlen, C.A., Chan, E.K., 2011. Induction of cytoplasmic rods and rings structures by inhibition of the CTP and GTP synthetic pathway in mammalian cells. PLoS ONE 6, e29690.

Carman, G.M., Henry, S.A., 1989. Phospholipid biosynthesis in yeast. Annu Rev Biochem 58, 635–669.

Chakraborty, K., Hurlbert, R., 1961. Role of glutamine in the biosynthesis of cytidine nucleotides in Escherichia coli. Biochimica et biophysica acta 47, 607–609.

Cheftel, J.C., 1995. High-pressure, microbial inactivation and food preservation. Food science and technology international 1, 75–90.

Chen, K., Zhang, J., Tastan, O.Y., Deussen, Z.A., Siswick, M.Y., Liu, J.L., 2011. Glutamine analogs promote cytoophidium assembly in human and Drosophila cells. J Genet Genomics 38, 391–402.

Chong, L., 2001. Molecular Cloning. Science 292, 446–446.

Daumann, M., Hickl, D., Zimmer, D., DeTar, R.A., Kunz, H.H., Mohlmann, T., 2018. Characterization of filament-forming CTP synthases from Arabidopsis thaliana. Plant J 96, 316–328.

Diekmann, Y., Pereira-Leal, J.B., 2013. Evolution of intracellular compartmentalization. Biochemical journal 449, 319–331.

Duggin, I.G., Aylett, C.H., Walsh, J.C., Michie, K.A., Wang, Q., Turnbull, L., Dawson, E.M., Harry, E.J., Whitchurch, C.B., Amos, L.A., 2015. CetZ tubulin-like proteins control archaeal cell shape. Nature 519, 362.

Dyall-Smith, M., 2008. The Halohandbook: protocols for haloarchaeal genetics. Haloarchaeal Genetics Laboratory, Melbourne 14.

Eichler, J., 2003. Facing extremes: archaeal surface-layer (glyco) proteins. Microbiology 149, 3347–3351.

Enache, M., Popescu, G., Dumitru, L., Kamekura, M., 2009. The effect of Na+/Mg2+ ratio on the amylase activity of haloarchaea isolated from Techirghiol lake, Romania, a low salt environment. growth (M) 1, 1–5.2.

Endrizzi, J.A., Kim, H., Anderson, P.M., Baldwin, E.P., 2004. Crystal structure of Escherichia coli cytidine triphosphate synthetase, a nucleotide-regulated glutamine amidotransferase/ATP-dependent amidoligase fusion protein and homologue of anticancer and antiparasitic drug targets. Biochemistry 43, 6447–6463.

Endrizzi, J.A., Kim, H., Anderson, P.M., Baldwin, E.P., 2005. Mechanisms of product feedback regulation and drug resistance in cytidine triphosphate synthetases from the structure of a CTP-inhibited complex. Biochemistry 44, 13491–13499.

Fukuchi, S., Yoshimune, K., Wakayama, M., Moriguchi, M., Nishikawa, K., 2003. Unique amino acid composition of proteins in halophilic bacteria. Journal of molecular biology 327, 347–357.

Garrity, G.M., 2012. Bergey’s manual of systematic bacteriology: Volume one: the Archaea and the deeply branching and phototrophic bacteria. Springer Science & Business Media.

Han, J., Lu, Q., Zhou, L., Liu, H., Xiang, H., 2009. Identification of the polyhydroxyalkanoate (PHA)-specific acetoacetyl coenzyme A reductase among multiple FabG paralogs in Haloarcula hispanica and reconstruction of the PHA biosynthetic pathway in Haloferax volcanii. Appl Environ Microbiol 75, 6168–6175.

Ingerson-Mahar, M., Briegel, A., Werner, J.N., Jensen, G.J., Gitai, Z., 2010. The metabolic enzyme CTP synthase forms cytoskeletal filaments. Nat Cell Biol 12, 739–746.

Juez, G., Rodriguez-Valera, F., Ventosa, A., Kushner, D.J., 1986. Haloarcula hispanica spec. nov. and Haloferax gibbonsii spec, nov., two new species of extremely halophilic archaebacteria. Systematic and Applied Microbiology 8, 75–79.

Kandler, O., König, H., 1998. Cell wall polymers in Archaea (Archaebacteria). Cellular and Molecular Life Sciences CMLS 54, 305–308.

Koshland Jr, D., Levitzki, A., 1974. 16. CTP Synthetase and Related Enzymes, The Enzymes. Elsevier, pp. 539–559.

Levitzki, A., Koshland, D.E., Jr., 1972. Role of an allosteric effector. Guanosine triphosphate activation in cytosine triphosphate synthetase. Biochemistry 11, 241–246.

Levitzki, A., Koshland Jr, D., 1971. Cytidine triphosphate synthetase. Covalent intermediates and mechanisms of action. Biochemistry 10, 3365–3371.

Lieberman, I., 1956. Enzymatic amination of uridine triphosphate to cytidine triphosphate. J Biol Chem 222, 765–775.

Liu, H., Han, J., Liu, X., Zhou, J., Xiang, H., 2011a. Development of pyrF-based gene knockout systems for genome-wide manipulation of the archaea Haloferax mediterranei and Haloarcula hispanica. J Genet Genomics 38, 261–269.

Liu, H., Wu, Z., Li, M., Zhang, F., Zheng, H., Han, J., Liu, J., Zhou, J., Wang, S., Xiang, H., 2011b. Complete genome sequence of Haloarcula hispanica, a Model Haloarchaeon for studying genetics, metabolism, and virus-host interaction. J Bacteriol 193, 6086–6087.

Liu, J.-L., 2010. Intracellular compartmentation of CTP synthase in Drosophila. Journal of Genetics and Genomics 37, 281–296.

Liu, J.L., 2016. The Cytoophidium and Its Kind: Filamentation and Compartmentation of Metabolic Enzymes. Annu Rev Cell Dev Biol 32, 349–372.

Long, C., Koshland, D.E., Jr., 1978. Cytidine triphosphate synthetase. Methods Enzymol 51, 79–83.

Long, C.W., Pardee, A.B., 1967. Cytidine triphosphate synthetase of Escherichia coli B. I. Purification and kinetics. J Biol Chem 242, 4715–4721.

Lynch, E.M., Hicks, D.R., Shepherd, M., Endrizzi, J.A., Maker, A., Hansen, J.M., Barry, R.M., Gitai, Z., Baldwin, E.P., Kollman, J.M., 2017. Human CTP synthase filament structure reveals the active enzyme conformation. Nat Struct Mol Biol 24, 507–514.

Lynch, E.M., Kollman, J.M., 2019a. Coupled structural transitions enable highly cooperative regulation of human CTPS2 filaments. bioRxiv, 770594.

Lynch, E.M., Kollman, J.M., 2019b. Coupled structural transitions enable highly cooperative regulation of human CTPS2 filaments. Nature Structural & Molecular Biology, 1–7.

Madern, D., Ebel, C., Zaccai, G., 2000. Halophilic adaptation of enzymes. Extremophiles 4, 91–98.

Noree, C., Sato, B.K., Broyer, R.M., Wilhelm, J.E., 2010. Identification of novel filament-forming proteins in Saccharomyces cerevisiae and Drosophila melanogaster. J Cell Biol 190, 541–551.

Ostrander, D.B., O’Brien, D.J., Gorman, J.A., Carman, G.M., 1998. Effect of CTP synthetase regulation by CTP on phospholipid synthesis in Saccharomyces cerevisiae. Journal of Biological Chemistry 273, 18992–19001.

Saitou, N., Nei, M., 1987. The neighbor-joining method: a new method for reconstructing phylogenetic trees. Molecular biology and evolution 4, 406–425.

Schmidt, I., Pfeifer, F., 2013. Use of GFP-GvpE fusions to quantify the GvpD-mediated reduction of the transcriptional activator GvpE in haloarchaea. Archives of microbiology 195, 403–412.

Stevenson, A., Cray, J.A., Williams, J.P., Santos, R., Sahay, R., Neuenkirchen, N., McClure, C.D., Grant, I.R., Houghton, J.D., Quinn, J.P., 2015. Is there a common water-activity limit for the three domains of life? The ISME journal 9, 1333.

Sun, Z., Liu, J.-L., 2019. Forming cytoophidia prolongs the half-life of CTP synthase. Cell Discovery 5, 32.

Thombre, R.S., Shinde, V.D., Oke, R.S., Dhar, S.K., Shouche, Y.S., 2016. Biology and survival of extremely halophilic archaeon Haloarcula marismortui RR12 isolated from Mumbai salterns, India in response to salinity stress. Scientific reports 6, 25642.

Willemoes, M., 2004. Competition between ammonia derived from internal glutamine hydrolysis and hydroxylamine present in the solution for incorporation into UTP as catalysed by Lactococcus lactis CTP synthase. Archives of biochemistry and biophysics 424, 105–111.

Woese, C.R., Kandler, O., Wheelis, M.L., 1990. Towards a natural system of organisms: proposal for the domains Archaea, Bacteria, and Eucarya. Proceedings of the National Academy of Sciences 87, 4576–4579.

Zhang, J., Hulme, L., Liu, J.L., 2014. Asymmetric inheritance of cytoophidia in Schizosaccharomyces pombe. Biol Open 3, 1092–1097.

Zhou, X., Guo, C.-J., Hu, H., Zhong, J., Sun, Q., Liu, D., Zhou, S., Chang, C.C., Liu, J.-L., 2019. Drosophila CTP synthase can form distinct substrate-and product-bound filaments. Journal of Genetics and Genomics.

