## Supporting Information for "CTP synthase forms cytoophidia in archaea"

### Table S1-S3

### Figure S1-S8

**Table S1. The amino acid alignment results.**

| Alignments | PYRG_THET8 | PYRG1_HUMAN | PYRG_SACS2 | PYRG_DROME | PYRG_ECOLI | PYRG_MYCTU | URA7_SACC | URA8_SACC |
|---|---|---|---|---|---|---|---|---|
| **Identities** | 53% | 51% | 50% | 49% | 49% | 50% | 48% | 46% |
| **Similarities** | 70% | 67% | 70% | 67% | 67% | 64% | 66% | 65% |

**Table S2. Strains and plasmids used in this study.**

| Strain or plasmids | Relevant characteristic(s) | Sources or references |
|---|---|---|
| **Strains** | | |
| *H. hispanica* DF60 | pyrF deletion mutant of *H. hispanica* | (Liu et al., 2011) |
| HhCTPS-GFP | GFP knocks into HhCTPS at C terminus of DF60 (pop-in) | This study |
| DH5α | Engineered *E.coli* cell with maximized transformation efficiency | This study |
| Transetta (DE3) | Derivative strain of BL21 with transformation efficiency and protein expression capability | TRANSGEN BIOTECH |
| HhCTPS-GFP-DE3 | Heterogeneous expression of HhCTPS tagged GFP in DE3 | This study |
| EcCTPS-GFP-DE3 | Expression of EcCTPS tagged GFP in DE3 | This study |
| **Plasmids** | | |
| pHAR | 4.0 kb; integration vector containing pyrFHh and its native promoter | (Liu et al., 2011) |
| pHAR-HhCTPS-C-GFP | Plasmid for construction of GFP to knock into C terminus of HhCTPS | This study |
| pWL502 | Expression vector containing pyrF and its native promoter | (Cai et al., 2012) |
| pWL502-GFP | Expression plasmid for GFP tag | This study |
| pWL502-HhCTPS-GFP | Expression plasmid for GFP tag fused to C terminus of HhCTPS | This study |
| pET28a | IPTG-inducible expression vector with His tag | This study |
| pET28a- HhCTPS-mCherry | Expression plasmid for mCherry tag fused to C terminus of HhCTPS | This study |
| pET28a-EcCTPS-mCherry | Expression plasmid for mCherry tag fused to C terminus of EcCTPS | This study |

**Table S3. Primers used in this study.**

| Primers | Sequences |
|---|---|
| HhCTPS-C-UF | CAGGTCGACTCTAGAGGATCCGTCAACTCCGAGAAGATG |
| HhCTPS-C-UR | GTGGCTCACCTCCTCAGTC |
| HhCTPS-C-GFP-F | GACTGAGGAGGTGAGCCACGCAGGATCCGCAGCAAGTAAAGGAGAAGAAC |
| HhCTPS-C-GFP-R | CGTCGACGTTCACCATCATTTGTATAGTTCATCC |
| HhCTPS-C-DF | TGATGGTGAACGTCGACGAGTTC |
| HhCTPS-C-DR | GAATCAGTTCCGCTAAGGTACCCCTCGACGACGTGGTTGG |
| HhCTPS-N-UF | CAGGTCGACTCTAGAGGATCCGAACTGGGACAGGCCGTC |
| HhCTPS-N-UR | ACCGACGGTGAACCCCGC |
| HhCTPS-NUCD-F | CGCGGGGTTCACCGTCGGTTGGTGAACGTCGACGAGTTC |
| PHAR-ID-F | CTCCGGTGACGCGTTCT |
| PHAR-ID-R | ATTACGCCAGATATCAAATTAATAC |
| HhCTPS-ID-F | GCTATCAAATCCGGCGTGCTC |
| HhCTPS-ID-R | GTCTCGAACTCCCGGATGAAC |
| HhCTPS-GFP-F | CAACACCGAGTTAGGAGATGGGATCCTGAATGCCGACCGAACCCGAAAC |
| HhCTPS-GFP-R | GGGAACCGCACACAAGAAAACGGTACCTCATTTGTATAGTTCATCCAT |
| PWL502-ID-F | ATTGGGCCGGAGATTGCACAGC |
| PWL502-ID-R | TCTCGATGCGGTCCTGAAGGTC |
| ECCTPS-F | CTTTAAGAAGGAGATATACCATGACAACGAACTATATTTTTG |
| ECCTPS-R | CATCGAGCCACCGCCACCCTTCGCCTGACGTTTCTGG |
| mCherry-F | GGGTGGCGGTGGCTCGATGGTGAGCAAGGGCGAGGAG |
| mCherry-R | CTTCCTTTCGGGCTTTGTTACTTGTACAGCTCGTCCATG |
| LOOP-PET28a-F | TAACAAAGCCCGAAAGGAAG |
| LOOP-PET28a-R | GGTATATCTCCTTCTTAAAG |
| HhCTPS-F | TTAAGAAGGAGATATACCATGCCGACCGAACCCGAAAC |
| HhCTPS-R | CATCGAGCCACCGCCACCGTGGCTCACCTCCTCAGTCG |
| PET28a-ID-F | TATAGGGGAATTGTGAGCGG |
| PET28a-ID-R | ACCCCTCAAGACCCGTTT |

### Figures S1-S8

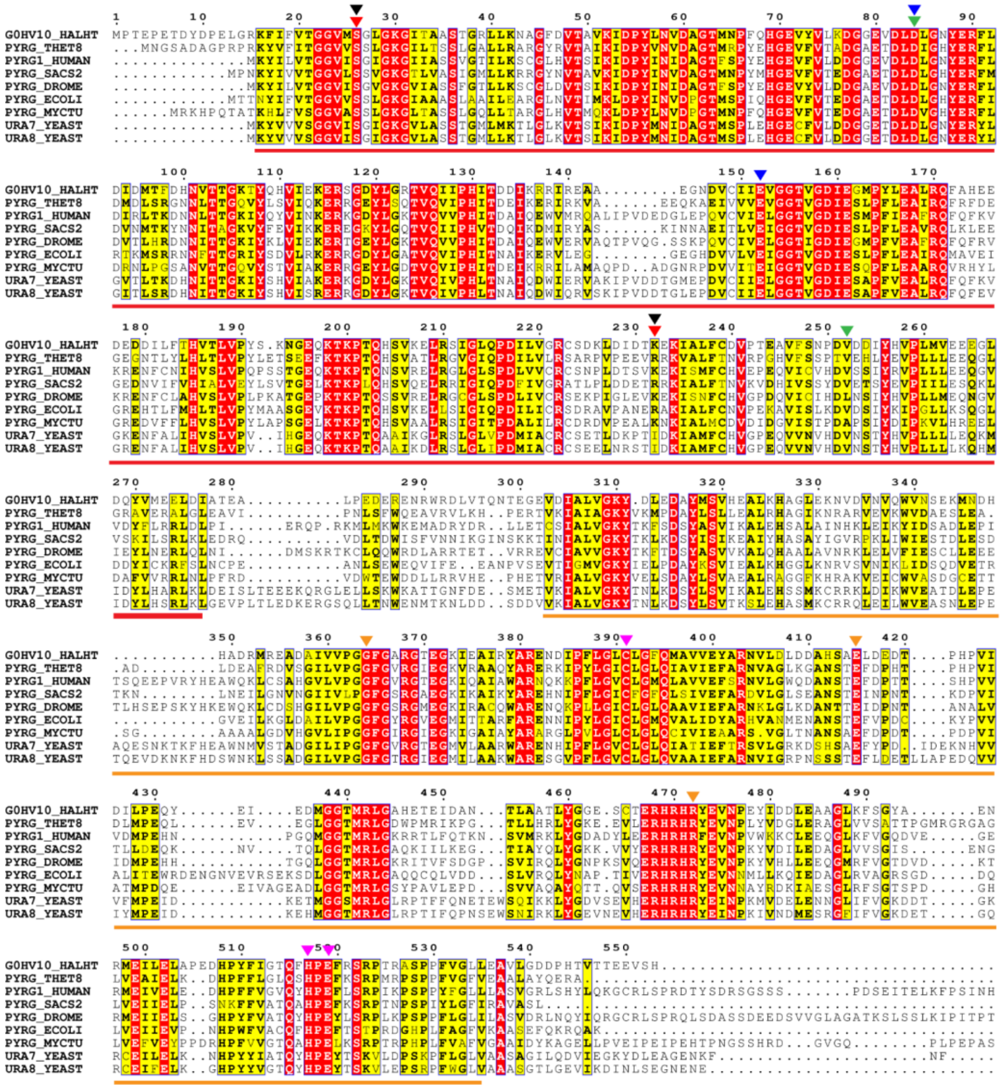

**Figure S1. The amino acid sequence alignment of CTPS in different species.** Alignment of amino acid sequences of *H. hispanica* CTPS (G0HV10_HALHT, WP_044951697.1) – homologous proteins from seven species (*Thermus thermophiles* [PYRG_THET8, WP_011228700.1], *Homo sapiens* [PYRG1_HUMAN, NP_001896.2], *Saccharolobus solfataricus* [PYRG_SACS2, WP_009990444.1], *Drosophila melanogaster* [PYRG_DROME NP_001287067.1], *E. coli* [PYRG_ECOLI, WP_000210878.1], *Mycobacterium tuberculosis* [PYRG_MYCTU, NP_216215.1], *Saccharomyces cerevisiae* [URA7_SACC, NP_009514.1; URA8_SACC, NP_012637.4]). Identical residues are shaded in red, and similar residues are shaded in yellow. Reverse triangles show some conserved residues: CTP binding site (red), UTP binding site (black), ATP binding site (green), metal binding site (blue), glutamine binding site (yellow), catalytic triad (purple). N-terminus ligase domain is marked by red line. Type Ⅰ glutamine amidotransferase (GATase 1) domain is marked by yellow line.

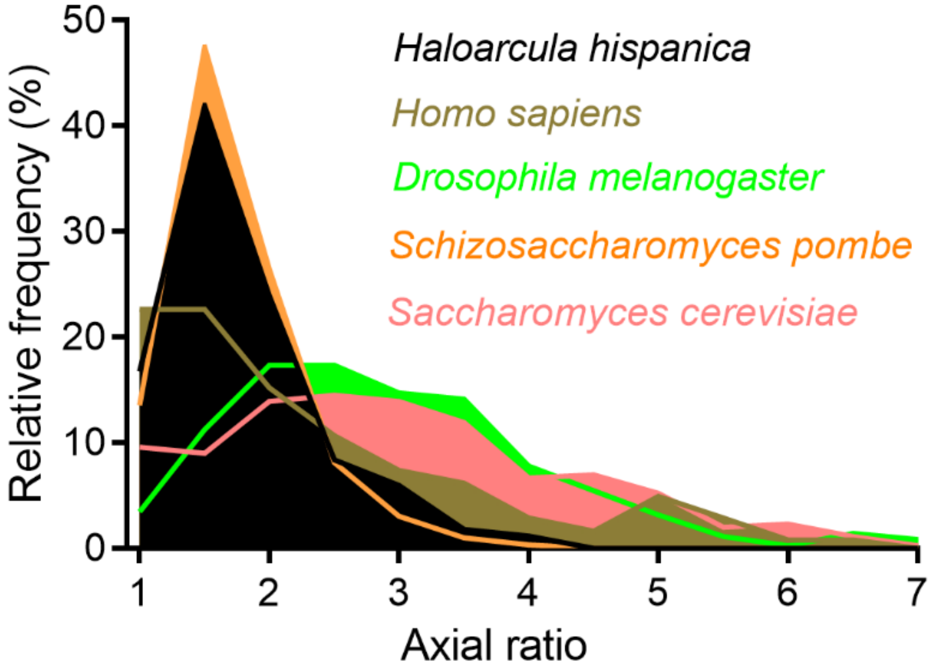


**Figure S2. Quantification of axial ratio of cytoophidia from different species.** The endogenous cytoophidia from *Haloarcula hispanica* are STED images; the others are confocal images. All images analysed by Fiji. Axial ratio = Major axis / Minor axis. n = 167 cytoophidia for *H. hispanica*, 243 cytoophidia for *Homo sapiens* (SW480), 346 cytoophidia for *Drosophila melanogaster* (ovary), 295 cytoophidia for *Schizosaccharomyces pombe*, 344 cytoophidia for *Saccharomyces cerevisiae*.

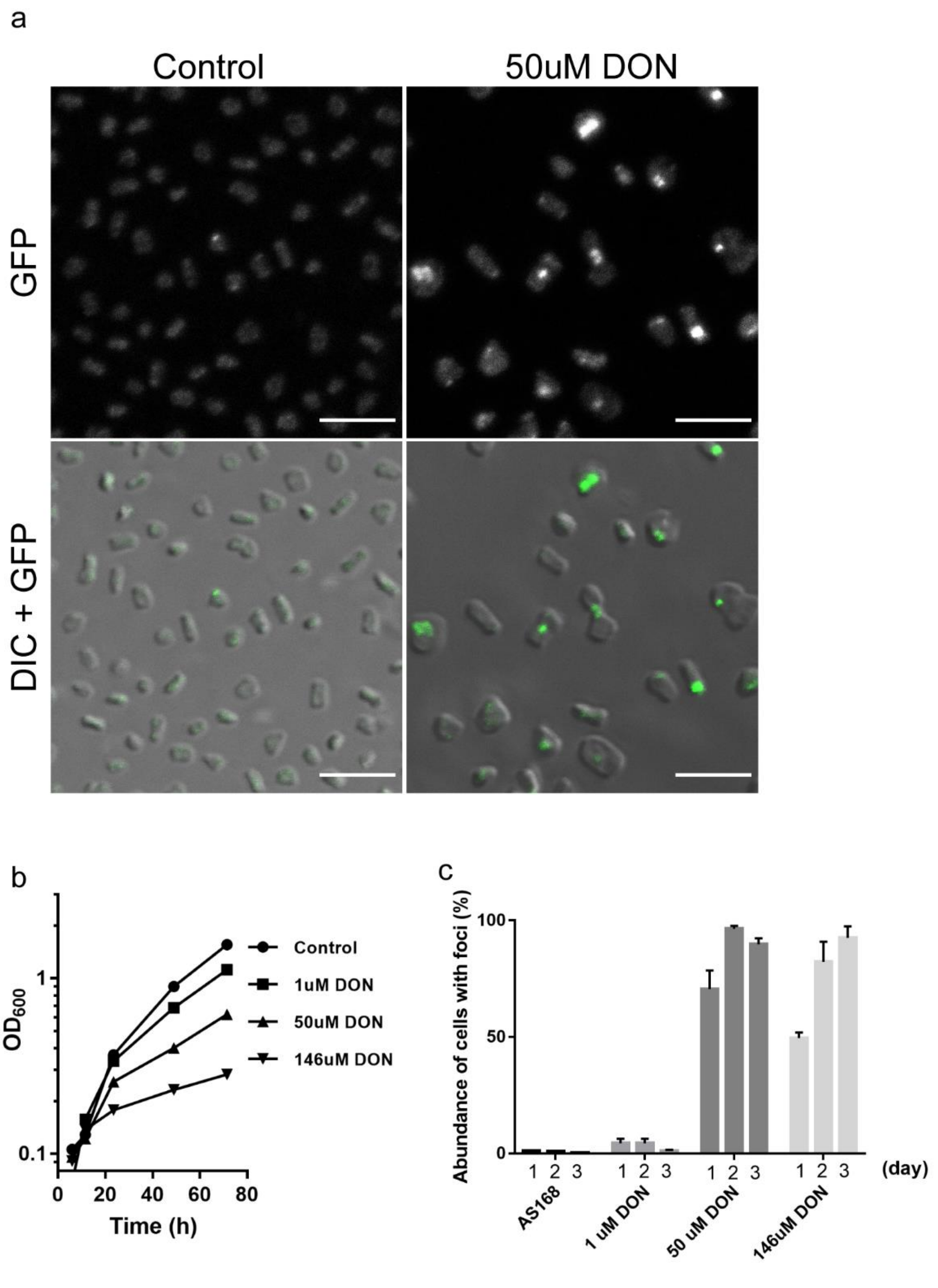


**Figure S3. A glutamine analog promotes the compartmentation of CTPS in *H. hispanica*. a**, After treating with 50 μM 6-diazo-5-oxo-l-norleucine (DON), a glutamine analog, for 3 days, HhCTPS-GFP shows a high proportion of cells with obvious foci. **b**, Growth curve of *H. hispanica* DF60 cultures treated with different concentrations of DON, with a log scale for y axis. **c**, DON treatment increases the dot signals. Cells were treated with 0 μM DON (n = 1117 for 1 day, 2538 for 2 days, and 3069 for 3 days), 1 μM DON (n = 1293 for 1 day, 7794 for 2 days, and 1372 for 3 days), 50 μM DON (n = 1539 for 1 day, 1097 for 2 days, and 2391 for 3 days), 146 μM DON (n = 1224 for 1 day,1302 for 2 days, and 1233 for 3 days). Mean ± SD. Scale bars, 5 μm.

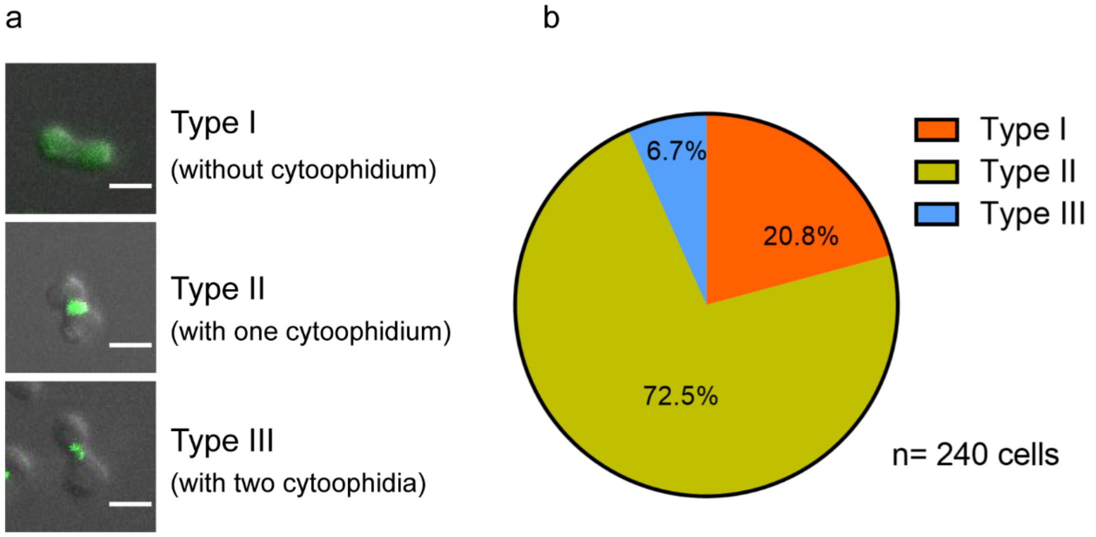


**Figure S4. The location of HhCTPS cytoophidia in the dividing cells treated by DON. a**, Representative images of three types of location of HhCTPS cytoophidia. Scale bar, 2 μm. **b**, Quantification of these three types.

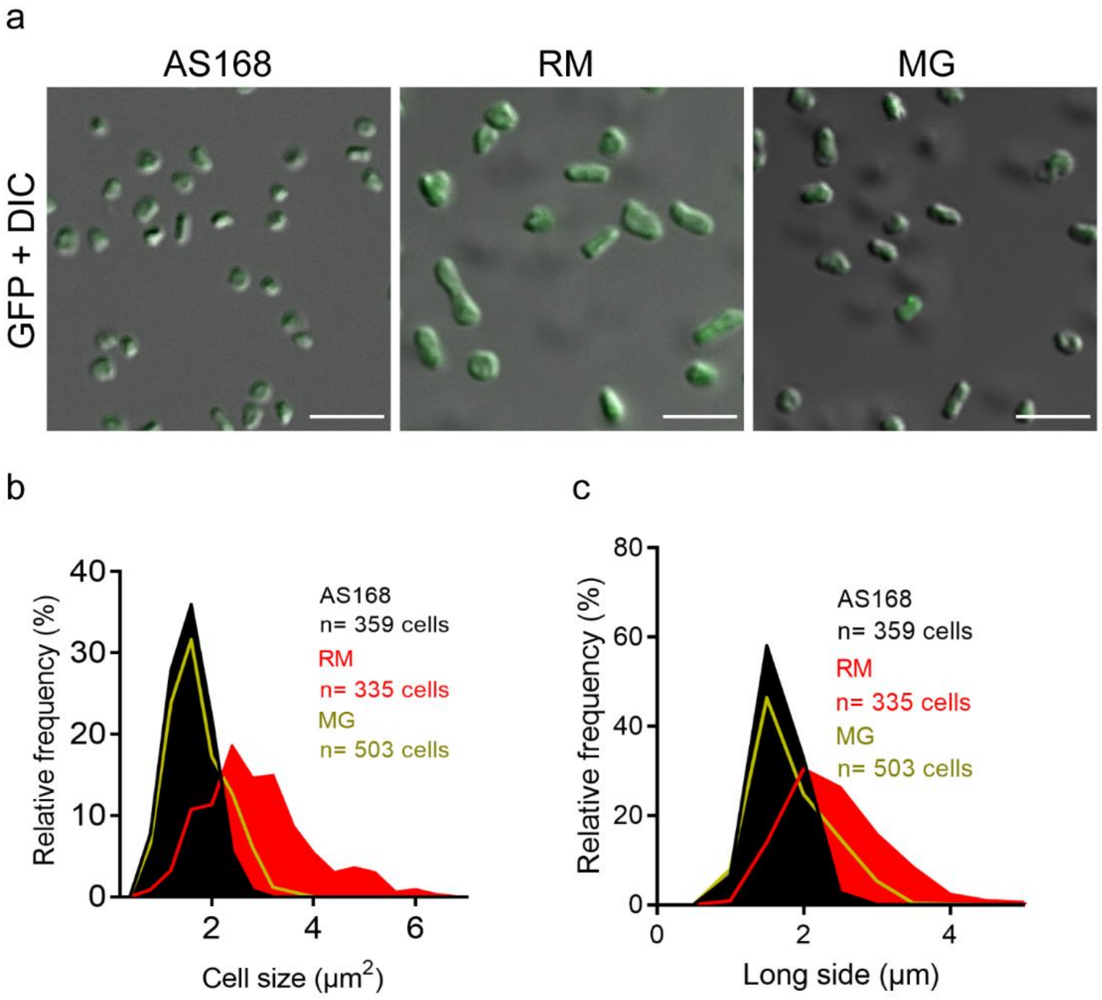


**Figure S5. Large cells are induced by resuscitation medium (RM). a**, Confocol images of HhCTPS-GFP cultured in AS168 medium, resuscitation medium (RM), nutrient-limited minimal medium (MG). **b**, **c**, Images were analyzed for cell size (**c**) and cell length (**d**). Scale bars, 5 μm.

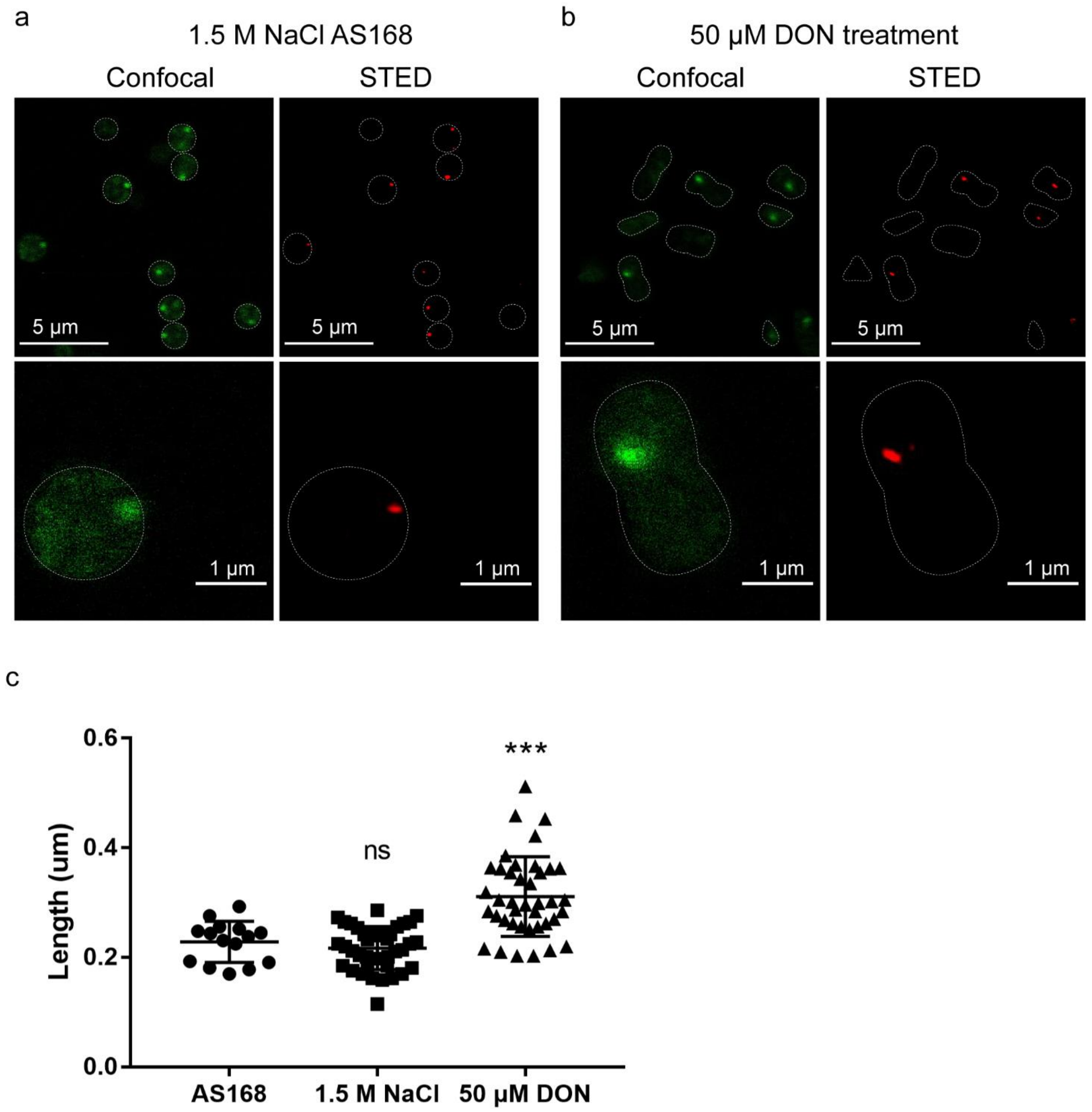


**Figure S6. Super-resolution images show elongated CTPS structures under low salinity and DON treatment. a**, **b**, STED images of HhCTPS-GFP in low salinity medium (**a**) and DON-supplemented medium (**b**). **c**, Quantification of length of elongated compartmentalized structure. n =15 in normal conditions, 39 in low salinity medium, and 40 under 50 μM DON treatment. Scale bars, 5 μm in upper pane of (**a, b**), 1 μm in lower pane of (**a, b**).

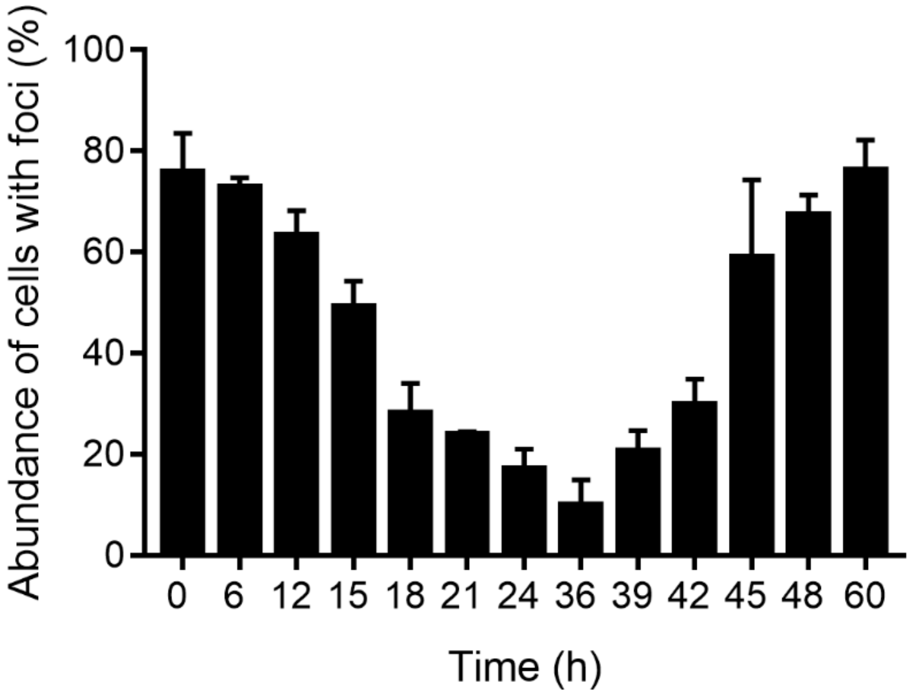


**Figure S7. The change in abundance of cytoophidia in OE-HhCTPSc GFP during the first 2.5 days of culture.** n= 987 cells for 0h (the start time point of culturing), 663 cells for 6h, 1095 cells for 12h, 1328 cells for 15h, 1365 cells for 18h, 1214 cells for 21h,1565 cells for 24h, 1329 cells for 36h, 1882 cells for 39h, 2666 cells for 42h, 1435 cells for 45h, 1908 cells for 48h, 855 cells for 60h. Mean ± SD.

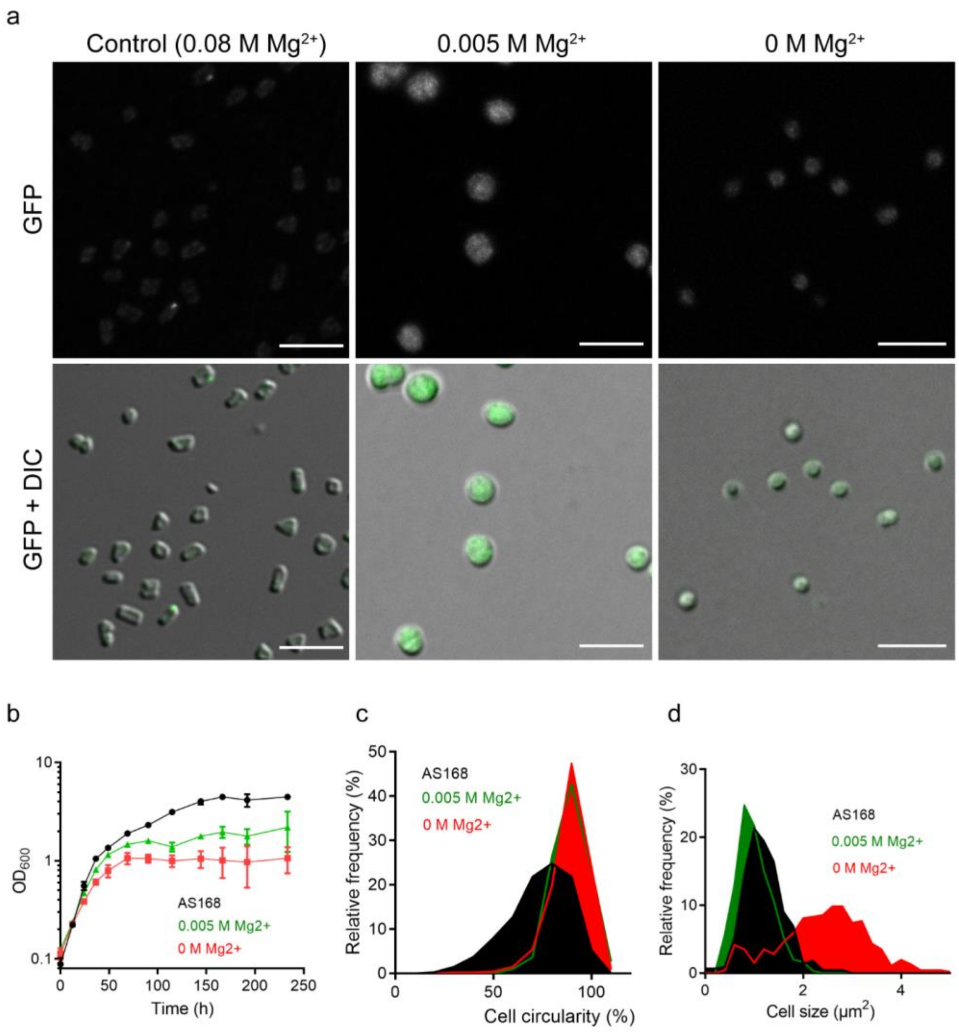


**Figure S8. *Haloarcula hispanica* tends to form a round cell shape in magnesium ion deprived or reduced AS168. a**, Confocol images of HhCTPS-GFP cultured in AS168 (0.08M $Mg^{2+}$), 0M $Mg^{2+}$ AS168, 0.005M $Mg^{2+}$ AS168 for 3 days. **b**, Growth curve of *H. hispanica* DF60 with a log scale for y axis. **c**, **d**, Images were analyzed for cell circularity (**c**) and size (**d**). n = 432 cells in AS168, 457 cells in 0M $Mg^{2+}$ AS168, and 300 cells in 0.005M $Mg^{2+}$ AS168. Mean ± SD. Scale bars, 5 μm.

### References

Cai, S., Cai, L., Liu, H., Liu, X., Han, J., Zhou, J., Xiang, H., 2012. Identification of the haloarchaeal phasin (PhaP) that functions in polyhydroxyalkanoate accumulation and granule formation in Haloferax mediterranei. Appl Environ Microbiol 78, 1946-1952.

Liu, H., Han, J., Liu, X., Zhou, J., Xiang, H., 2011. Development of pyrF-based gene knockout systems for genome-wide manipulation of the archaea Haloferax mediterranei and Haloarcula hispanica. J Genet Genomics 38, 261-269.
